# Contact-dependent Communication Shapes the Mesoscale Spatial Organization of Bacterial Swarms

**DOI:** 10.64898/2026.05.31.726327

**Authors:** E. E. Garling, M. Byers, E. Bradley, J. D. Meiss, K. A. Gibbs

## Abstract

Scientists use microscopy images to visualize and describe organisms and their interactions. Microscale visual inspections of individual cells and colonies, paired with genetic approaches, have revealed key developmental steps within microbial communities including, for example, swarms, biofilms, and fruiting bodies. However, there is a lack of formal, quantitative descriptions of these structures. This gap limits understanding of the structure and relevance of cell clusters, especially regarding development and interactions. Here, we develop biologically informed mathematical methods to formally assess cell clusters that assemble and dissolve over time in bacterial colonies. Our approaches to this analysis of the developmental stages of swarming, a type of collective migration, focus on cell length (as a proxy for developmental stage), cell-to-cell contacts, and groupings. We apply these to data from *Proteus mirabilis* strains with genetic disruptions in different aspects of its communication mechanisms to explore how identity signaling and cell-to-cell contact affect population structure at different scales. We found that kin recognition and contact-dependent, cell-cell communication govern population architecture during collective migration, such as swarming. This integration of microbiology with applied mathematics and computer science is a frontier for analyzing experimental data, leading to testable biological insights.

**Significance:** Collective behaviors are often exhibited by social organisms. Open questions regarding these behaviors include: how do communication and familial identity influence local interactions, and how do local interactions combine to create collective behaviors? Qualitative descriptions from visual examination of microbes can be useful in approaching these questions, but formal quantification of these structures will be essential if we are to understand the mechanisms that underlie organismal behavior and organization. This manuscript bridges microbiology and applied mathematics, interleaving the mathematics and the experiments in an iterative fashion. We offer critical advances to better understand, analyze, and characterize the multiscale structures that form and dissolve as a colony engages in collective migration. Our findings suggest that cell-cell communication plays an important role in population architecture, specifically that communication may “prime” cells for collective migration and prevent stagnation during life stage transitions.

## A. Introduction

Community structure relies on interactions between individuals, whether animals, fungi, or bacteria. In honey bees, for instance, clusters form due to scenting behaviors involving the exchange of pheromones through the air (1). Contact-dependent communication is one example of bacterial interactions; kin recognition is one of its modifiers (2–4). In microbes, community structure is critical for colonization in diseases, fitness in mixed populations, and metabolism. Since Antonie van Leeuwenhoek, microbiologists have used increasingly higher magnifications to examine the activities of individual microorganisms, i.e., the microscale. However, a critical gap blocks forward progress in understanding individual cells in the context of microbial community structures: formal analysis of groups of cells, i.e., structure at the *mesoscale*. Closing this gap would allow one to answer longstanding questions—such as the role of cell-cell communication or kin identity in community structure and maintenance—which remain unanswered in multiple microbial systems.

This barrier has blocked progress on mesoscale analysis. Traditional techniques for image analysis often struggle to quantitatively assess cell connectedness and the spatial organization and architecture of bacterial communities (5–7). In addition, human visual bias can distort the interpretation of microscopy images (8). Recent research has overcome some of these challenges in the context of populations with minimal to no motility (9–11), populations with cells that are uniform in shape (12, 13), and labeled subpopulations (14, 15). However, many microbial populations contain cells of varying lengths and shapes that are motile and function collectively. To the best of our knowledge, broadly applicable mathematical tools for mesoscale analysis of complex microbial communities have not yet been developed.

We move beyond these limitations using a combination of formal mathematical methods and targeted experiments. In Section (B), we discuss the opportunistic pathogenic bacterium, *Proteus mirabilis*, studied in this paper, which utilizes contact-dependent kin identity as a regulator of collective migration (16). *P. mirabilis* cells move collectively as a group, called a “swarm,” on hard surfaces (17, 18). This migration is dependent on a developmental cycle in which cells oscillate between being short (< 4 µm) and nearly uniformly rod-shaped to becoming elongated and snake-like (up to 40+ µm long) (19–22). For wild-type populations, these distinct life stages can be reliably measured through quantitative changes in cell length distributions within a heterogeneous population (23). Kin recognition is active during this swarm development cycle; the current model is that kin identity acts either before or after collective migration to block access to colony benefits, such as shared molecules, nutrients, or predator protection, by non-kin cells (24). Using a genetics approach that selectively modifies the communication mechanisms, combined with cell length analysis and novel mathematical analyses that focus on cell adjacency and the topology of cell groupings, we address the longstanding question: how does kin recognition influence the population’s architecture?

To the eye, microscopy images of swarms of non-kin cells appear to have structures that are distinct from the swarms of kin (25, 26), but quantitative analysis to verify this distinction has been previously unavailable. Mathematical approaches from the fields of computational geometry and computational topology offer a promising path to formalize such qualitative observations, thereby delivering meaningful, quantifiable descriptions of the structures in complex images of polymorphic cell populations (see e.g., Figure 1A). In Section (C), we introduce mathematical methods for formally characterizing the degree of cell-cell contact and the sizes of cell groupings in *P. mirabilis* colonies. Together, these methods provide an interaction network that quantifies the potential for communication among adjacent cells. Using these methods, alongside genetic modifications to communication, we show that cell-cell communication and kin identity alter how neighbors interact and influence the structures that form in swarms. Our methods differ from existing order parameters used in the study of biological aggregation (e.g., packing fraction (27) or average nearest-neighbor distance (28)). Indeed, the generality of the mathematical methods described in this paper allows us to account for a collection of differently shaped individuals, and our results reveal patterns—and biological variability—across experiments. Our goal is to formally characterize bacterial complexity, self-organization, immense morphological changes, collective dispersion, and coalescence.

**Figure 1.**
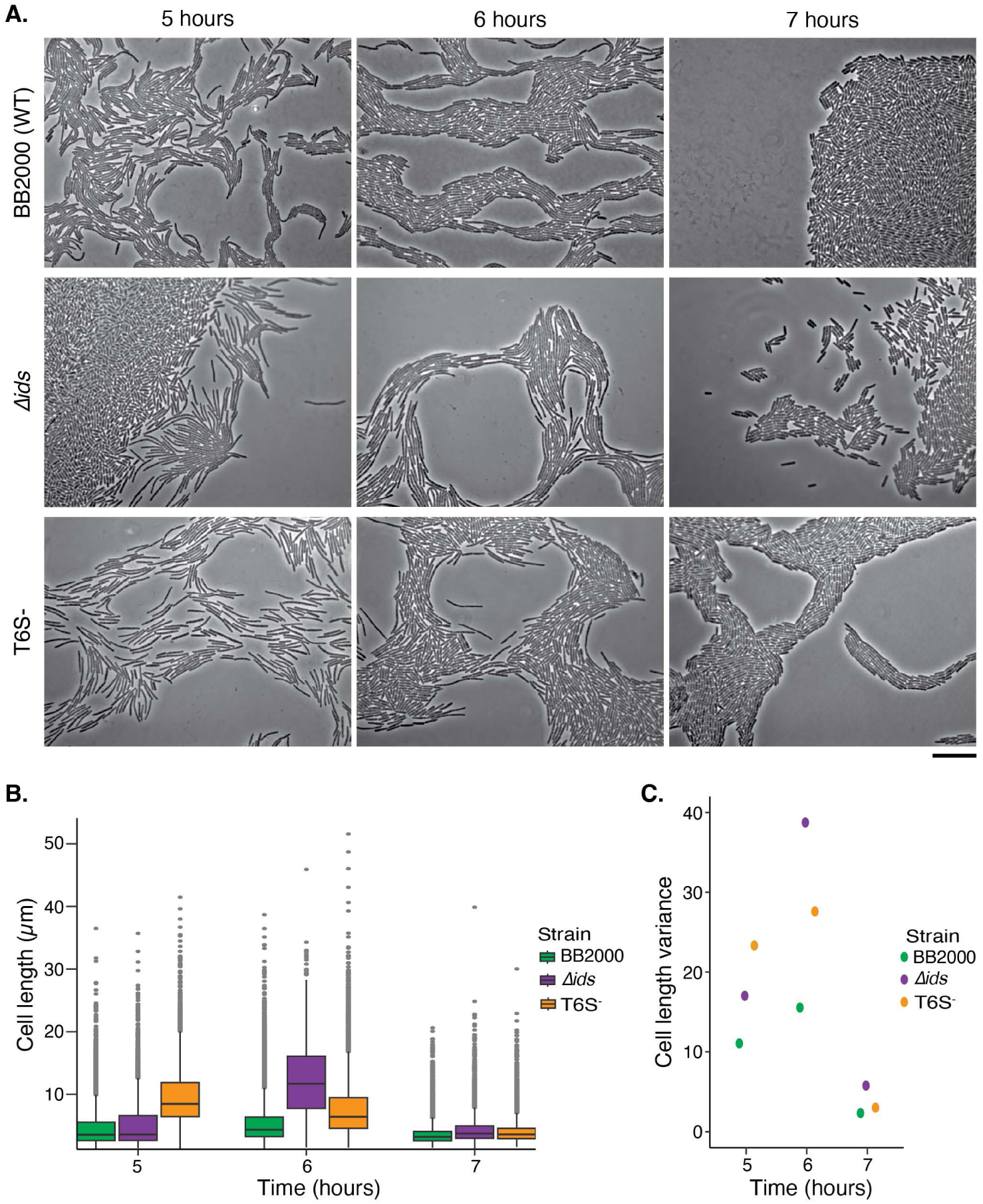
Disruptions in intercellular communication alter cell physiology and population structures. (A) Representative phase contrast images, at three timepoints, of swarms from wild-type strain BB2000 (WT) and two derived mutant strains with a disruption in non-lethal identity signaling (*Δids*) or in the cell-cell communication machinery (T6S-). Images are swarm initiation at five hours, collective swarming at six hours, and consolidation at seven hours. Scale bar: 10 µm. (B) *P. mirabilis* cell lengths computed using Swarmetrics for the populations as labelled. Only cells present in a single layer were analyzed. Each graph is a boxplot where the box represents the 1^st^-3^rd^ quartile, and the middle line is the median. Single points above and below the box show the points outside these quartiles. This morphological analysis of the BB2000 data set was previously published (23) and is provided for comparison to the mutant strains. (C) Variance of the cell lengths as a swarm development measure. The largest variances occur during collective swarming; the smallest during consolidation.

## B. Biological Methods

### Strains and media

The wild-type *P. mirabilis* strain is BB2000 (29), from which the two mutant strains are derived: *Δids* (30) and T6S- [*P. mirabilis* BB2000 *tssB*_T95G_, which was constructed and confirmed as described previously for CCS05 using the pKNG101-derived vector, pCS34 (25)]. *P. mirabilis* is stored at -80°C and daily maintained on LSW^-^ agar at 4°C (29). For propagation, cells are grown overnight aerobically in LB broth (Lennox) at 37°C. CM55 blood agar base (Thermo Fisher Scientific, Waltham, MA) is the media for swarming.

### Microscopy

Overnight cultures are diluted and normalized in LB broth (Lennox) to an optical density at 600 nm of 1.0 and then inoculated onto a CM55 agar pad on a microscope slide as previously described (30). Microscopy is performed as previously described (25). Briefly, the leading edge of the swarms and inoculum are imaged using a Leica DM5500B (Leica Microsystems, Buffalo Grove, IL) and a CoolSnap HQ2 cooled charge-coupled-device (CCD) camera (Photometrics, Tucson, AZ). A new microscope slide is used for each timepoint and biological replicate; all are inoculated at the same time in parallel. MetaMorph version 7.8.0.0 (Molecular Devices, Sunnyvale, CA) is used for image acquisition. Images analyzed for the mathematical methods are available in the supplementary materials.

### Image analysis

To avoid obfuscating downstream analysis, images are first cropped, if necessary, to remove layered cells, oversaturated regions, or areas of excess debris. The perimeter of the image is excluded, since it often includes cells that are partially out of the frame. Cell masks and length values are acquired using the Swarmetrics pipeline, which is described in Garling et al. (23). In the current Swarmetrics pipeline, approximately 10% of cells in each microscopy image are missed, so the contact networks between the cells are likely an underestimate, particularly at the edges of the image where neighboring cells are cut off. The *Δids* strain is particularly difficult to segment and includes images with >10% of unaccounted cells. This rate of detection is comparable to other bacterial detection algorithms (5). The resulting data is exported as .csv files for the analysis described in Section (C). See GitHub <u>repository</u> <https://github.com/mbyers31/biomorphological-analysis-suite> for the methods described herein.

## C. Results and Mathematical Methods

Kin recognition through direct cell-to-cell communication appears to alter cell development—specifically, morphology or behaviors. For example, disruptions in cell-cell communication of self-identity reduce the width of *P. mirabilis* swarm colonies by as much as 80% (25), specifically through the transfer of Ids self-identity proteins via a contact-dependent Type IV secretion system (T6SS) (31, 32). Comparative analyses of individual cells have not yet been pursued, though by eye, the communication-disrupted mutant strains appear to have different lengths and form atypical cell groupings (Figure 1A). In this study, we analyze three phases of the *P. mirabilis* swarm development cycle using data gathered via phase contrast light microscopy. Using multiple independent isolates, we consider the potential role of cell-cell communication of kin identity in swarm structure. Figure 1A shows representative images of wild-type strain BB2000 (WT) and its derived mutant strains: *Δids*, which is completely disrupted in the communication of its non-lethal Ids self-identity but not in all communication (30), and T6S-, which completely lacks T6SS-dependent cell-cell communication (33, 34). These strains are shown at three sequential stages of swarm development: swarm initiation (entry in movement, hour 5), collective swarming (fastest movement, hour 6), and consolidation (cells stop moving and some divide, hour 7) (22, 23). Interestingly, in the microscopy images, the structures of the groupings in the imaged populations vary between strains and across time.

For comparative analysis of the differences, we first need to assess whether each strain is at the same development stage, i.e., progressing similarly to wildtype. This analysis also examines how communication and kin identity impact cell morphology. We measure each cell’s length using Swarmetrics, an image capture and analysis pipeline (23), resulting in a final data set of 55,000 cells over all samples (Supplemental Table 1). For wildtype, both median cell lengths and their variances rise at hour 6 and then fall to their lowest values at hour 7 (Figures 1B, 1C), as previously reported (23). These results show that wild-type cells begin to elongate during swarm initiation (hour 5), becoming (as a population) longest during collective swarming (hour 6), and shorten during consolidation (hour 7), likely due to cell division. Cell lengths for the *Δids* population follow the same pattern, except that the *Δids* cells are, on average, longer than wildtype during collective swarming (Figure 1B). By contrast, the T6S- population has longer cells than wildtype during both swarm initiation and collective swarming, but the distribution of short cells is like those for both wildtype and the *Δids* strain during consolidation (Figure 1B). The variances for cell length within each mutant strain’s population follow the same pattern as wildtype, with the greatest values during collective swarming and the lowest during consolidation; cells have the largest spread of cell lengths at hour 6 (Figure 1C). These data suggest that these three strains are all undergoing swarm development at roughly equivalent time periods, allowing for comparative analysis across strains.

An important aspect of the cell-length data discussed above is the ratio of short to long cells, which is a marker of cell development. We previously found that long cells dominate populations of *P. mirabilis* during collective swarming, while short cells are dominant at the swarm initiation and consolidation stages (23). To further explore this issue, we partitioned the cells in the current experiments into two groups, using a 4 µm cutoff—double the length of a non-swarming cell. The percentages of the populations in these two groups show that wildtype and the *Δids* strain share a pattern in which the longer cells predominate during collective swarming (Supplemental Table 2). By contrast, the T6S- population has essentially the same fraction of long cells during swarm initiation and collective swarming (Supplemental Table 2). Short cells dominate during consolidation for all strains, although less so for the *Δids* strain.

Altogether, this data shows that morphological changes in cells occur when communication is disrupted. However, a challenge for this type of aggregate analysis of the cell lengths is that the data are spatially decontextualized. Figure 1 tells us about the overall size distributions but does not capture the geometry of the cell groupings that appear in the micrographs of Figure 1A. *In other words, we are missing some critical information about the mesoscale.* Formal, geometric assessment of this kind of structural information, we hypothesize, is key to analyzing and understanding the differences between the temporal evolution of the colony structure of these strains.

To quantify the mesoscale structure that is present in the micrographs (Figure 2A), we introduce a measure that we call **δ-adjacency**, which is designed to quantify cell-cell contact, a critical feature in communication. Direct physical contact is needed for the T6SS-dependent transfer of identity proteins like Ids (31, 32). In addition, efficient swarm migration requires cell-cell contact, as the bacterial flagella are thought to intertwine and allow for greater force propulsion (20, 35). To formally quantify this property, we define the δ-adjacency for each cell in a scatterplot (Figure 2B) to be the *fraction of its perimeter that is within a distance δ of some other cell*. The rationale for the choice of δ in this metric stems from image processing. The software in our pipeline creates cell masks that do not account for the bulkiness of intertwined flagella, thereby creating apparent gaps in the image between neighboring cells that are actually in physical contact. The parameter δ accounts for this shortcoming: cells are considered to be adjacent only if they are separated by a gap of at most δ. The value we choose, δ = 0.59 µm, is the average of the distance between pairs of visually observed neighboring cells across several images (Supplement Figure 1). We use this value in all adjacency calculations reported in this paper.

**Figure 2.**
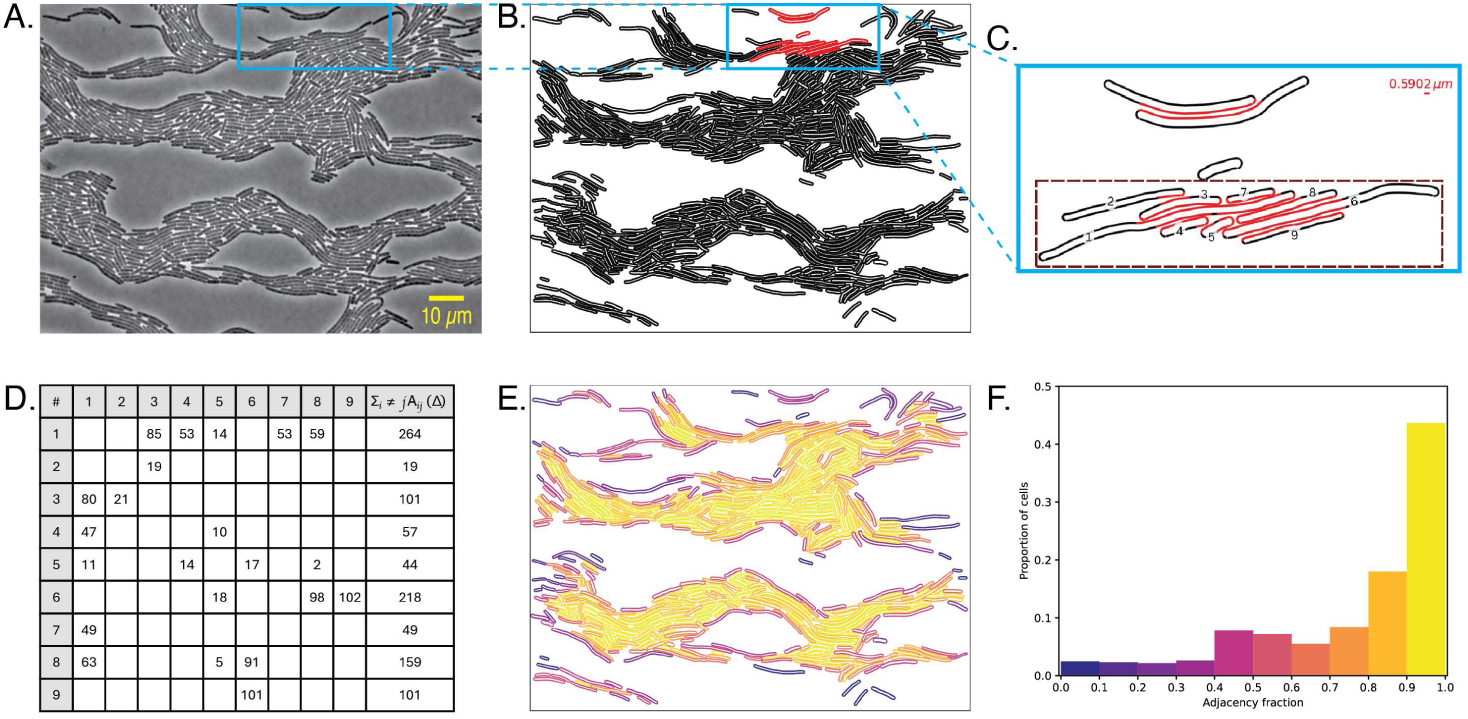
Adjacency metric for quantifying cell-to-cell contact. (A) Phase contrast image of wildtype at 5 hours. (B) Scatter plot of the cells in Panel A. (C) Magnified image of the red-highlighted cells from Panel B. Points on each cell’s perimeter that are within δ = 0.59 µm of any other cell are shown in red. The δ parameter allows us to identify contact in the face of imaging artifacts. (D) The δ-adjacency values are stored in a matrix whose *i,j*^th^ entry represents the number of perimeter points of cell *i* that are δ-adjacent to cell *j*. The *i*^th^ row sum of this matrix represents the total number of perimeter points of cell *i* that are δ-adjacent to *any* other cell in the image. (E) Heatmap of the δ-adjacency fractions for the scatter plot in Panel B. (F) Histogram of adjacency fractions for panel E.

To compute the δ-adjacency value for a cell, we consider each sample point along its perimeter to determine how many are within δ of another cell. When there are *N* cells in an image, the result is an *NxN* matrix, *A*(δ), the **δ-adjacency matrix**. Here each element, *A_ij_*(δ), is the number of points on the perimeter of cell *i* that are within δ of cell *j.* In this matrix, there is one row and one column for each cell; the rows represent “outgoing” contact (who each cell is touching) and the columns represent “incoming” contact (who is touching each cell). Figure 2C demonstrates the results of this computation for the small patch of the scatterplot shown in Figure 2B, with δ-adjacent points highlighted in red. In this example, 47 of the perimeter points of cell 4 are δ-adjacent to cell 1 and 10 to cell 5, so *A*_4,1_(δ) = 47 and *A*_4,5_(δ) = 10. Figure 2D shows a portion of the δ-adjacency matrix corresponding to the nine cells in the highlighted region of Figure 2C. For the sake of readability, entries in the matrix that are zero (corresponding to pairs that are not δ-adjacent) are not shown. Note that this matrix will typically not be symmetric, since the different cells can have different sizes and different perimeter point spacing.

We can find each cell’s *aggregate* δ-adjacency by computing the row sums of the δ-adjacency matrix: Σ_j= 1…n_*A_ij_*(δ) — that is, the total number of each cell’s perimeter points that are within δ of *any* other cell. To allow for comparison between cells with different lengths, we normalize this quantity, dividing by the number of points along the perimeter of each cell, to give the **δ-adjacency fraction**, which runs from 0 to 1. The heatmap in Figure 2E demonstrates the results of this calculation on the scatterplot in Figure 2B, color-coding each cell according to its δ-adjacency fraction. Heatmaps like this can bring out important features of the colony structure that are not immediately apparent from a visual examination. For example, some of the regions in Figure 2E that appear equally dense to the eye in the micrograph (Figure 2A) and scatterplot (Figure 2B) have a higher δ-adjacency fraction than others, indicating different degrees of cell-cell contact. Since this information is given for each cell, it can be used to reveal differences between regions in the colony—as well as patterns that could relate to important biological properties such as “rafting,” where cells are “parallel” along their longest axis (20), which assumes a degree of contact. Such analysis goes beyond a qualitative description of scatter plots and provides an *explicit* measure of a cell’s opportunity to communicate and share resources.

We can further aggregate the information in a δ-adjacency heatmap in the form of a histogram by binning the δ-adjacency scores, as shown in Figure 2F. In this example, 43.7% of the cells have an adjacency score between 0.9 and 1.0: *i.e.*, 90-100% of their perimeter is adjacent to some other cell (encoded as yellow in Figure 2E and 2F). While this lumped analysis does lose the geometric information that is present in Figure 2E, such high-level information about the overall proportion of cells that are in close contact can be useful in identifying large-scale differences between different phases and strains The more cells with high δ-adjacency fraction that we see on the histogram, for example, the more opportunity there is for direct cell-to-cell communication across the population in the image, indicating possible benefits from group interactions. Histograms can then be aggregated across images so that consistent signals become visible above the natural variations of biological systems and experimental replicates.

The results of applying the δ-adjacency metric to the *P. mirabilis* data sets, introduced in Section (B), are shown in Figure 3A. These bring out information about the micro- and meso-scale structure that is not apparent from the cell-length data or the microscopy images (see also Supplemental Figures 2-7). The top row of Figure 3A shows that δ-adjacency in the wild-type strain increases as the population transitions from swarm initiation (hour 5) to the collective swarming phase (hour 6), where there is visible rafting, but then drops during consolidation (hour 7). This drop was unexpected, as consolidated cells visually appear to be densely packed, when viewed in comparison to hour 6; in other words, the δ-adjacency results indicate that visual perception of “dense packing” can be deceptive. The T6S- strain reaches its highest δ-adjacency during consolidation; the previous timepoints show a “pick-up sticks” organization with less cell-cell contact. For *Δids,* there is more variability between the three images, and the structure alternates between similarity to the wild-type and T6S- strains. Notably, the δ-adjacency for the *Δids* strain is much lower at hour 7 (consolidation) than the other strains. Similarities in the heatmaps are present across a single data set, yet the heat maps for the full series of images in the supplement show differences across replicates within a single time-point.

**Figure 3.**
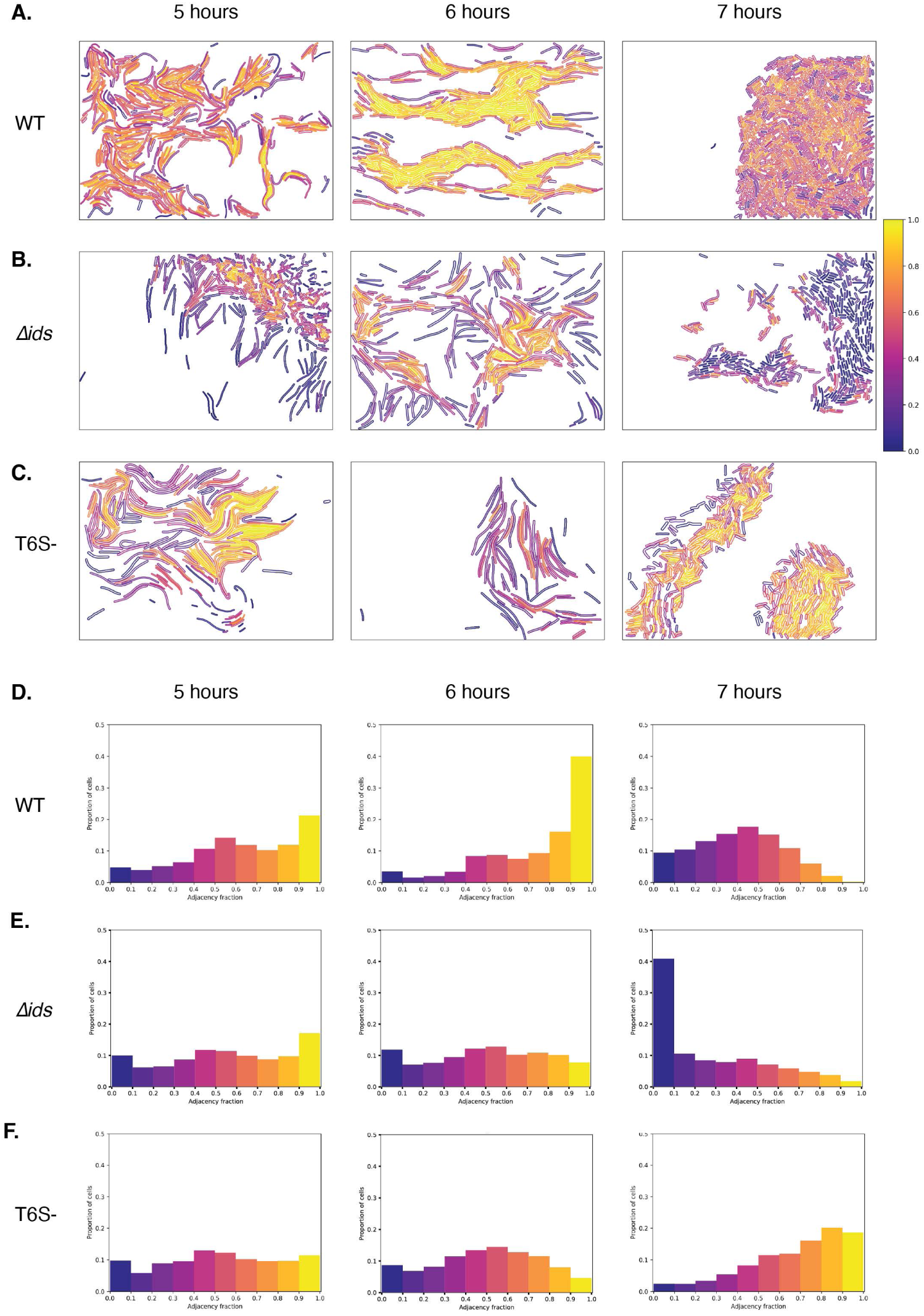
Adjacency reveals mesoscale structures. (A-C) Heatmaps of δ-adjacency for sample scatterplots for each strain at each timepoint. (D-F) Histograms of δ-adjacency fraction for all scatterplots for each strain.

To assess patterns in these results in the face of this variability, we use the δ-adjacency fraction histogram from Figure 2F to aggregate the results across all images for each strain and timepoint; see Figures 3D-F. Each histogram bin is the proportion of cells in all scatterplots of this strain at this timepoint that fall into the normalized adjacency score range specified on the horizontal axis. From these plots, we can make some interesting observations about differences between strains and across time. On average, the wild-type populations have higher adjacency at hour 5; this increases slightly at hour 6, with a notable decrease at hour 7. The *Δids* strain does not have a similar change in adjacency across the developmental phases, while the T6S- strain displays the inverse pattern from the wildtype: the highest adjacency values occur during consolidation, at hour 7. The adjacency distribution of the *Δids* populations remains remarkably similar for hours 5 and 6 but shows a notable reduction in δ-adjacency across all populations during consolidation, indicating that cells are in less direct contact with each other at hour 7. The T6S- population has a relatively flat adjacency distribution for hour 5, exhibits fewer cells with higher adjacencies during hour 6, and has a notable shift at hour 7, when most cells have more than half of their perimeter within δ of another cell. The T6S-populations exhibit the highest adjacency during consolidation.

Overall, it is striking that the fraction of a cell’s body that is in contact with others varies with the phase of the swarm development cycle. In view of the biological differences among these engineered strains, these results show that identity signaling and communication (via T6SS) contribute to the degree of cell-cell contact. Disruptions in cell-cell communication alter the proportional adjacency amounts and timing, as compared to wildtype. Note that the loss of communication (in the T6S- mutant strain) leads to less adjacency during both swarm initiation and active swarming as compared to wildtype, raising the specific question as to whether cell-cell communication and identity signaling work to help “prime” cells for collective swarming. To begin answering this question, we need a formal quantification of the mesoscale structures in the colony: specifically, of the sizes and shapes of the *groups* of cells that are in contact with one another. Based on the biology, isolated cells have no immediate opportunity for direct, physical (T6SS-dependent) cell-cell communication. In contrast, adjacent cells do have the potential for this form of communication (33, 36). Multi-cell contact may even reflect “network” potential for communication, similar to quorum sensing (37, 38) but with contact.

To formalize this, we use the information in the δ-adjacency matrix to identify groups of bacteria that are in contact with one another by building a graph that captures the clustering structure (Figure 4). This graph contains a vertex for each cell; a nonzero entry of A_ij_(δ) indicates pairwise adjacency between cells *i* and *j*, giving an edge between the corresponding vertices. We then analyze the clusters in this graph using the tools of topological data analysis (TDA) (39). In the TDA framework, points in a data set are defined to be **ε-connected** if they are within ε of each other and groups of ε-connected points are called ‘connected components.’ In our context, ε = δ, and we call a connected component an *island*: a set of cells, each of which is δ-adjacent to at least one other cell in the set. For example, the zoomed-in region in Figures 4A and 4B contains two islands: one containing nine adjacent bacteria and one containing two. The final cell in this region is a *singleton*, an isolated cell that has no opportunity for communication or resource sharing. (In topological data analysis, this would be referred to as a δ-isolated cell: a connected component of size one.) Figure 4C shows the islands and singletons in the scatterplot from Figure 2B, distinguishing them by color. The largest islands (orange and blue) contain 56.95% and 37.71% of the 655 cells, respectively; the smaller islands, shown in various other colors, each contain between 0.31% and 1.83% of the total. In addition, there are 11 singletons that together comprise 1.68% of the total cell count.

**Figure 4.**
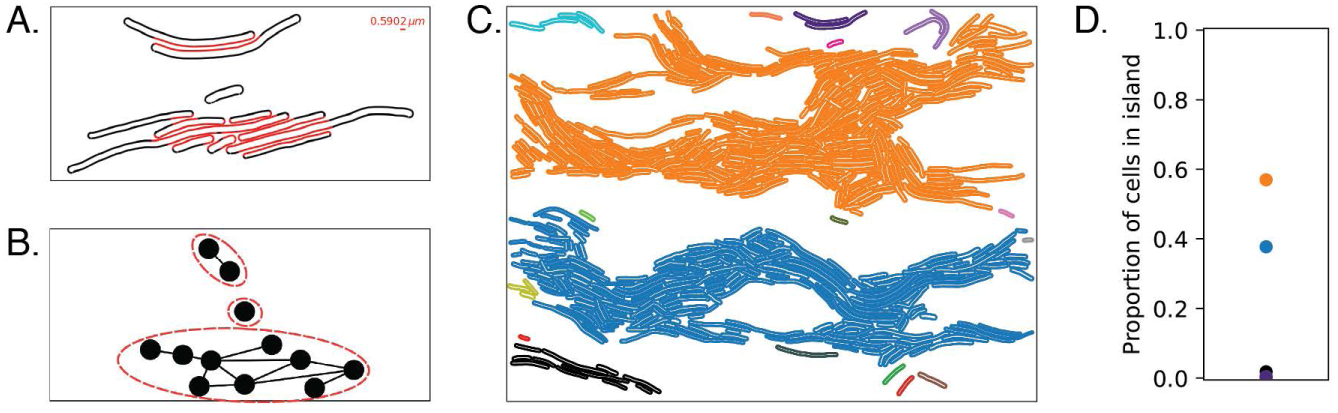
Building a graph from the δ-adjacency metric to identify cell groupings. (A) Duplicated from Figure 2C, this image shows δ-adjacent points highlighted in red. (B) The δ-adjacency graph for the 12-cells in panel A. Each vertex corresponds to a cell, and each edge connects a pair of cells that are δ-adjacent. A connected component of the graph—a group where all cells are δ-adjacent to at least one other cell in the group—is an *island*. There are two islands in this panel. The isolated cell between the islands is a *singleton*. (C) The island structure of the scatter plot of Figure 2. (D) Island size proportions for panel C, calculated as a fraction of the total number of cells.

To quantify this information, we make a plot that contains one point for each island, with the vertical position showing the proportion of cells that it contains (Figure 4D). We use proportion—rather than the number of cells—so that it is possible to compare images with different cell counts. Since these numbers are reported as proportions, they must sum to 1; thus, there can be at most one island with a proportion greater than 50%. We separately tabulate the number of singletons to capture the patterns in a series of experiments, making a box-and-whisker plot of the number of singletons in all the corresponding frames.

This way of analyzing cell groupings can be useful in elucidating important biological information. Figure 5 shows the islands and singletons for the images of Figure 3, using color to distinguish groups. The number and size of the islands varies both with time and with strain. At hour 6, all three strains have two predominant islands, but the shapes of each of these larger islands, as well as the distribution of smaller islands and singletons, are quite different. The wild-type images also contain more singletons at hour 6 than the other timepoints. The compositions of the cell groupings in the *Δids* and T6S- mutant strains are visibly different from each other and from the wild-type strain.

**Figure 5.**
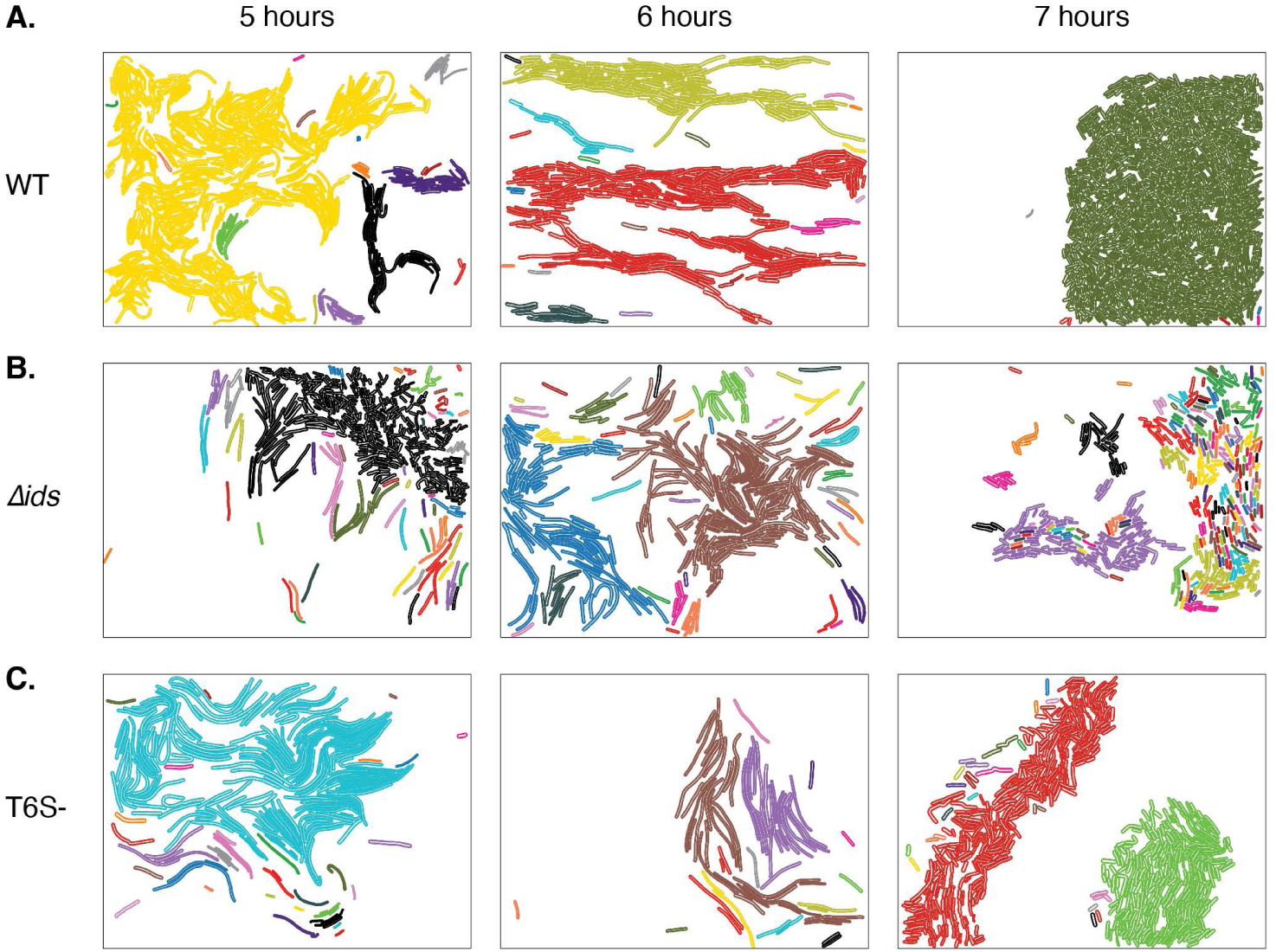
Identification of the islands and singletons for the scatterplots of Figure 3A, with each grouping shown in a different color. (A) Wildtype. (B) The *Δids* strain, which has partial communication. (C) The T6S- strain, which does not have T6S-dependent cell-cell communication.

While the images in Figure 5 are suggestive, they are not quantitative. To formalize the information that they contain, we use the size distributions introduced in Figure 4D to reveal and quantify patterns. These plots show island size proportions with a dot for each island, separating the image frames horizontally. These, as seen in Figures 6A-C, bring out both the patterns and the biological variability across the image frames for each condition (strain and timepoint). For example, at hours 5 and 6, there is a striking contrast between wild-type and the two mutant strains at the extremes of the size range: there are more large islands in the former and more small ones in the latter two. Indeed, this visualization reveals several interesting aspects of the mesoscale structure within a *P. mirabilis* strain.

**Figure 6.**
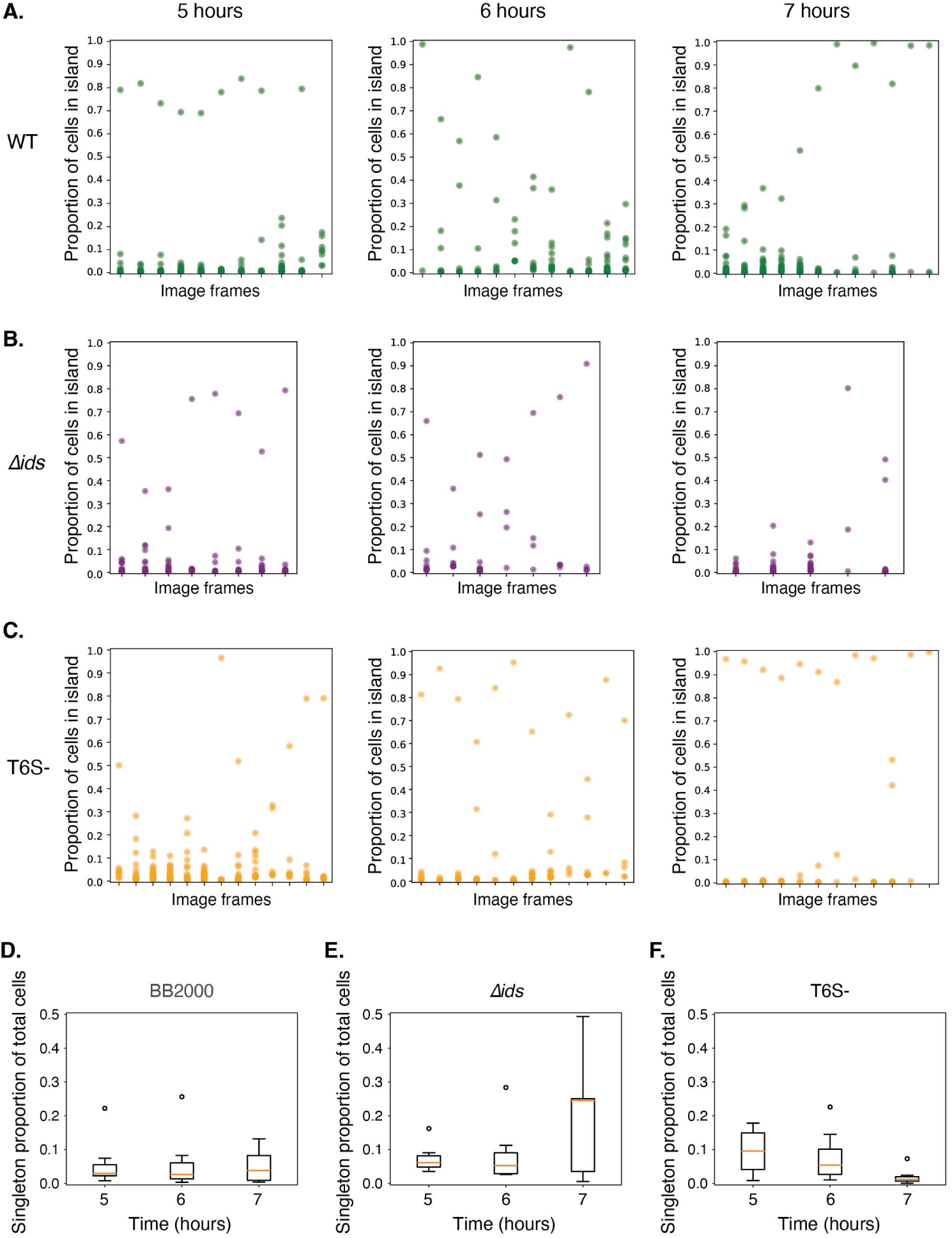
Analysis of the island sizes can formally quantify variations in the mesoscale structure between strains and frames at each timepoint. (A-C) The proportion of cells in each image that are contained in each island for each strain, image frame, and timepoint. (A) Wildtype. (B) The *Δids* strain. (C) The T6S- strain. (D) Box-and-whisker plots of the singletons for each of these conditions. Strain name is written above each plot.

Island size distributions vary significantly with time for wildtype (Figure 6A) and the T6S-mutant strain (Figure 6C). In contrast, the *Δids* mutant strain has similar proportions of islands for each timepoint. Specifically, for the wild-type population, the frames are dominated by a single large island containing ∼70–80% of the cells at hour 5. There are comparatively fewer small islands (under ∼20%), and just two image frames contain only disconnected small islands (Figure 6A). At hour 6, there is no single, dominant island for most of the image frames, suggesting that collective swarming leads to the “melting” or dispersal of these large islands, producing a distribution of island sizes that spans the full range of possibilities. By hour 7, there are two dominant regimes for island structure: a single, very large island containing 80–90% of the cells, with fewer small islands (like hour 5 but with bigger islands) or—for about half of the image frames—a spectrum of medium-size islands. This indicates that the mesoscale structure of the colony is not rigid, but rather that there is a continuum of possibilities.

These results support a hypothesis that contact-dependent communication plays a role in island formation: the island sizes are different when communication is disrupted. The *Δids* population shows little difference in island structure at the three timepoints: no single pattern in the distributions among image frames is visible (Figure 6B). Note, however, that large islands (those containing over ∼85% of cells) are mostly absent from the *Δids* populations; this suggests that this mutant strain forms smaller connected communities than the wild-type strain.

The island structure of the T6S- mutant also shows patterns that are distinct from those of the wildtype (Figure 6C). At hour 5, its dominant distribution is a collection of small- and medium-sized islands, except for perhaps three exceptional image frames. As in wildtype and the *Δids* mutant strain, the island sizes vary considerably at hour 6, during collective swarming. However, at hour 7, all frames but one have a single, very large island containing more than 80% of the visible cells; the exception contains two islands of approximately 40% and 50% of the cells (almost 90% total). These data indicate that the T6S- mutant strain forms one large connected “mat” of cells during consolidation, to a larger degree than seen with wildtype.

Taken together, these results suggest that the complete loss of communication *reduces* cell-cell connection during swarm initiation, is not impactful during collective swarming, and *increases* it during consolidation. By contrast, the loss of partial (non-lethal) communication in the *Δids* strain disrupts island structure, resulting in distributions similar to collective swarming throughout the swarm development cycle.

These analyses of the larger islands tell only a portion of the story; the singleton data are also revealing. Figure 6D shows box-and-whisker plots of the proportion of singletons. For wildtype, singletons consistently represent a small percentage of the cells—often less than 10%. In other words, wild-type cells tend to form groups. In contrast, both mutant strains have a higher proportion of singletons at hours 5 and 6. The T6S- strain has virtually no singletons at hour 7, during consolidation; in this phase, cells are generally found in comparatively large islands. The *Δids* strain has a higher proportion of singletons at every timepoint than the wildtype—and the greatest proportion during consolidation—which is consistent with the reduced δ-adjacency (Figure 3E). Taken together with the island size distributions, these results show that disrupting cell-cell communication and kin identity impacts how cells maintain contact. This phenomenon would not happen without the involvement of the corresponding molecular mechanisms (and signaling outputs).

## D. Discussion

Many existing studies of microbial collectives focus on the actions of individuals (the microscale) or the outcomes of the population as a whole (the macroscale), omitting that of intermediate sized groups: the mesoscale. The mesoscale is not only a critical bridge between the extremes, but also a key player in the dynamics. To address this need, we focused on complementary approaches: cell length distributions and the geometry of cell groupings that form and dissolve during the developmental phases of a *P. mirabilis* colony. We developed and leveraged new methods to analyze the geometry of these structures, applying them to the results of targeted experiments to derive biological insights about cell-cell communication. We found that *P. mirabilis* colonies form cell groupings of different sizes and that patterns in these sizes vary over the stages of swarm development. Of particular note is our result that cell-cell communication—specifically the transfer of T6SS-dependent self-identity signals—regulates both the morphology of cells and their degree of interaction with neighboring cells.

These findings not only agree with the well-known prediction that genetic relatedness promotes cooperation (24, 40) but also support a new hypothesis about the mesoscale structures that organize the network of connected cells within a swarm. Other biological studies have shown that bacterial populations can have increased fitness (growth, survival) in social groups; for example, that larger groups have increased surfactant in biofilms, protecting them against threats like phage and antimicrobials, and that quorum sensing can contribute to one population’s dominance in mixed consortia (41–44). In *P. mirabilis* swarms, group size might increase access to nutrients through surface occupation or facilitate protection against stresses like phage. The data and analyses presented here, in context of the field, suggest a new biological model: kin recognition prompts populations to form highly adjacent, connected components (groups) that together form a *mesoscale* interaction network.

This application of computational mathematics to microbiology has moved the needle. While previous work, which relied largely on qualitative descriptions derived from visual examination of microbes is certainly useful, it is essential to have a *formal* quantification of these structures if we are to truly understand the mechanisms that underlie organismal behavior and organization. There have been some recent developments along these lines in the context of uniformly shaped microbial cells with no motility (7)—as well as a significant body of work on other types of biological aggregations (27, 45), where the individual agents can be approximated as points, circles, or ellipsoids. Our work is more general: our mathematics is designed to handle complex biological patterns and the variations that occur amongst experiments. Further, our experiments leverage these analyses to study the biological mechanisms underlying group behaviors. Our methods enable automated analysis of large data sets, which may permit spatial analysis to move from genetic composition (e.g., transcriptomics and metagenomics) to physical, organismal population dynamics, as in, for instance, in studies of animal microbiomes (46). The results of this combined mathematical/biological approach are objective and reproducible (overcoming natural experimental and biological variability), enabling validation of existing hypotheses about cell-cell communications, as well as the formation of new ones—which can in turn inform the design of new experiments.

A striking finding in this analysis is that visual density does not imply physical contact. Perceiving groups of cells as functionally organized or coordinated when they are not was first termed by (47) as the “clustering illusion.” Existing techniques for the analysis of biological aggregations, such as density and nearest-neighbor analysis (27, 28), cannot resolve such an illusion. Our tight focus on adjacency, however, allows us to reveal these visual density paradoxes quite clearly. For example, even though, to the human eye, there appears to be less “grouping” of T6S- cells at all phases, when compared to wildtype, this is an illusion, as we saw using the δ-adjacency analysis (recall Figure 6).

Our comparison of the wildtype, *Δids*, and T6S- mutant strains suggests that communication plays an important role in population architecture. The island analysis that we introduce in this paper—a new way to identify contact networks and thereby facilitate a specific understanding of the biologically-relevant structure of the swarm—is based on analysis of the connected components in a graph that captures the δ-adjacency relationships between cells in an image. The sizes of these groups bring out salient information about network structure in a novel way. For example, wild-type populations are highly dynamic, forming large islands during swarm initiation, which then disperse during active swarming and then re-form during consolidation, perhaps suggesting a sort of dynamic coordination amongst the cells. In contrast, cells lacking all communication (T6S-) fail to aggregate into islands during swarm initiation and form oversized “mats” during consolidation. This suggests that communication may “prime” the cells for collective migration and prevent stagnation during consolidation. Finally, cell groupings in the *Δids* populations exhibit almost no change over time. Instead, the population appears to be stuck in a sort of “permanent migration state” consisting of disorganized small and medium islands. We posit that non-lethal identity signaling in this case may act as a signal that guides the transition into different life phases and/or organizes cells into rafts during collective migration.

Many opportunities for future work remain. A first goal would be to develop methods for characterizing other important attributes of the mesoscale structure, such as alignment. Although the term “alignment” is intuitive, it is surprisingly difficult to formalize, especially for cells with complicated shapes. Alignment and adjacency are related—two cells with high relative δ-adjacency should be designated as aligned, regardless of their shapes—but what constitutes a sufficiently “high” value is not obvious. Using the adjacency fraction and direction for each edge in a δ-adjacency graph would result in a weighted and directed graph that could give a quantitative metric for this attribute. Another potentially interesting aspect of mesoscale structure relates to the holes that are visible in many *P. mirabilis* colonies. The ideas of *persistent homology,* another powerful tool from topological data analysis (48), could help quantify such structures. It could also be useful to vary the threshold at which an edge is added between two vertices of an adjacency graph to explore the effects on the structure of the cell-cell communication network.

Our mathematical methods should extend smoothly to other collections of elongated bacteria that engage in collective migration, such as *Vibrio parahaemolyticus* (49), many of which remain poorly understood due to restricted genetics, cultivation, and image analysis tools. Our analyses might also help understand the role that contact-mediated interactions between *Capnocytophaga gingivalis* cells play in multi-species communities (50). We are excited to apply our mesoscale-analysis techniques to systems that involve physical contact between heterogeneous agents, like those in biofilms, engineered soil crusts, or synthetic metabolic communities (51–53), though a needed first step is improvement in the underlying segmentation and detection algorithms. Finally, we are looking forward to using these new methods to study how kin recognition and cell-cell communication regulate community structures, using genetic manipulations to further dissect the intracellular and intercellular pathways.

The methods presented here mark a significant shift from qualitative microscopy to a quantitative, spatial analysis of microbial spatial ecologies. Formal characterization of the structures in a bacterial colony has many affordances: not just comparison and contrast, but real insights into biological mechanisms. Key to this is effective interleaving of mathematics and microbiology. We used data from laboratory experiments to develop, test, and refine the mathematical methods; these insights then led to new hypotheses and the design of laboratory experiments for their validation. This synergistic approach allowed us to go beyond the descriptive and led to insights about cell-cell communication and its role in the formation of multiscale structures that emerge during the developmental phases of the colony—insights that are relevant to network development, maintenance, and expansion.

## Acknowledgements

We would like to thank Achala Chittor for construction of the T6SS-disrupted strain as well as members of the Gibbs lab for thoughtful discussions and comments on the manuscript. We thank the following funders of this research: the Alfred P. Sloan Foundation (grant number G-2022-19509 to K.A.G.), the National Science Foundation (Graduate Research Fellowship to E.E.G.), the University of California, Berkeley, the University of Colorado Boulder, and the Santa Fe Institute. We especially thank the David and Lucile Packard Foundation, who nucleated this collaboration by fortuitously having K.A.G. and E.B. present neighboring posters at a meeting.

## Supplemental Figures

**Supplemental Figure 1.**
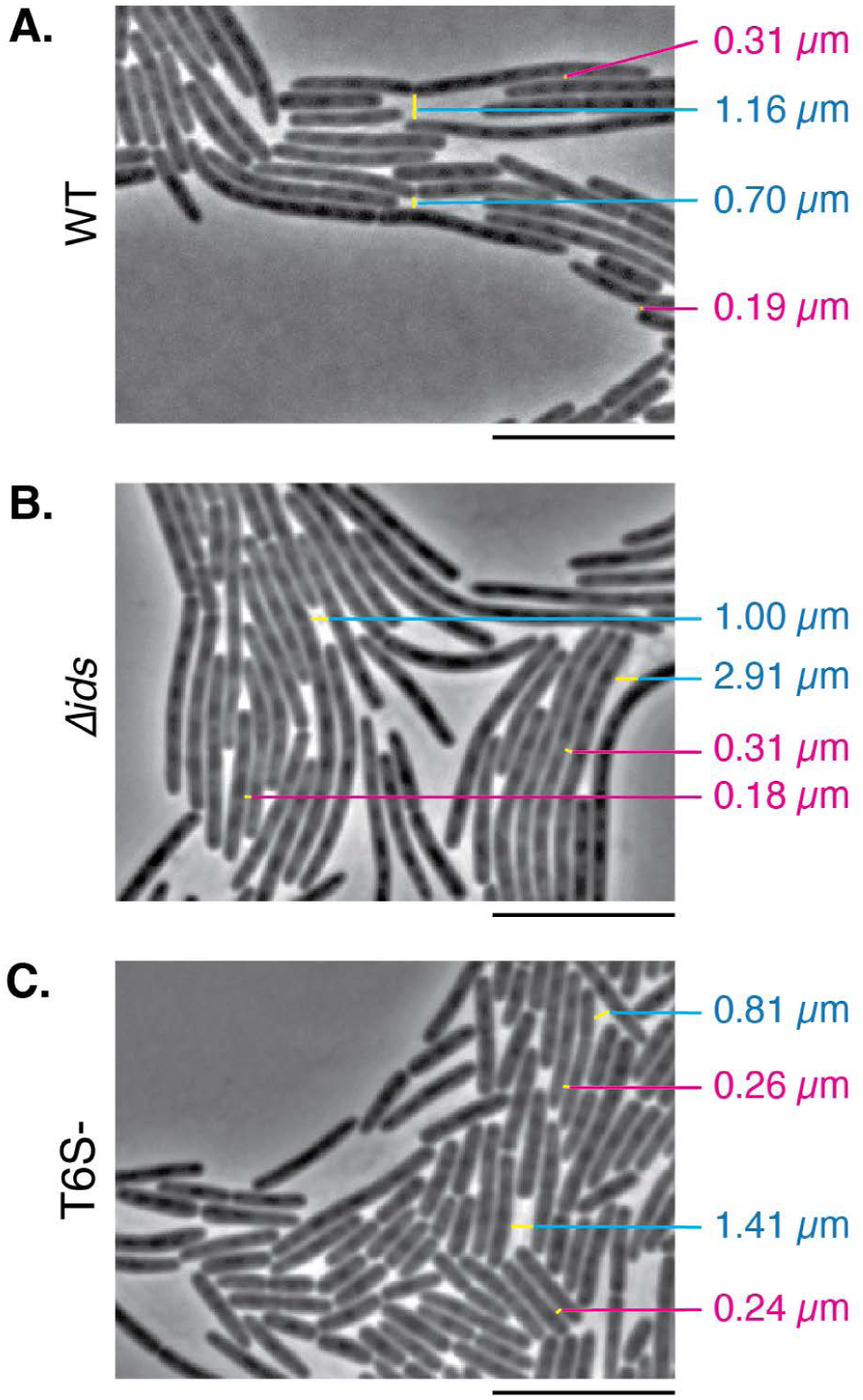
Measurement of δ-adjacency is based on calculations from phase contrast images. To determine the parameter δ, we measured distances between adjacent (magenta labels) and non-adjacent (cyan labels) cells, as labeled in demonstration on this figure. The value we choose, δ = 0.59 µm, is the average of the distance between pairs of visually observed neighboring cells across several images. (A) Wildtype. (B) *Δids*. (C) T6S-. Scale bar, 10 µm.

**Supplemental Figure 2.**
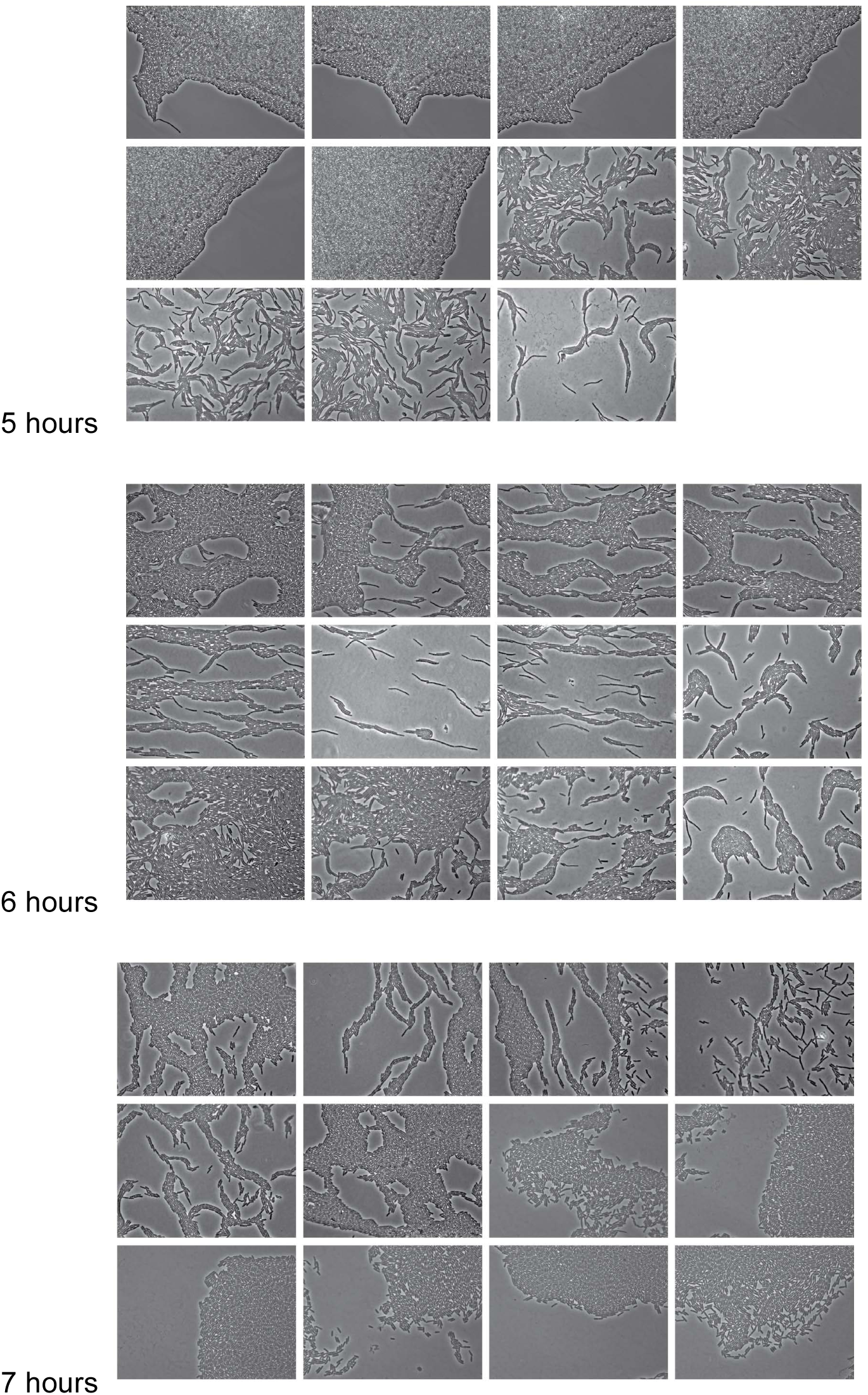
Phase contrast images of wild-type *P. mirabilis* strain BB2000 used for analysis. Each individual image is ∼ 126.4 µm wide by ∼ 94.4 µm tall.

**Supplemental Figure 3.**
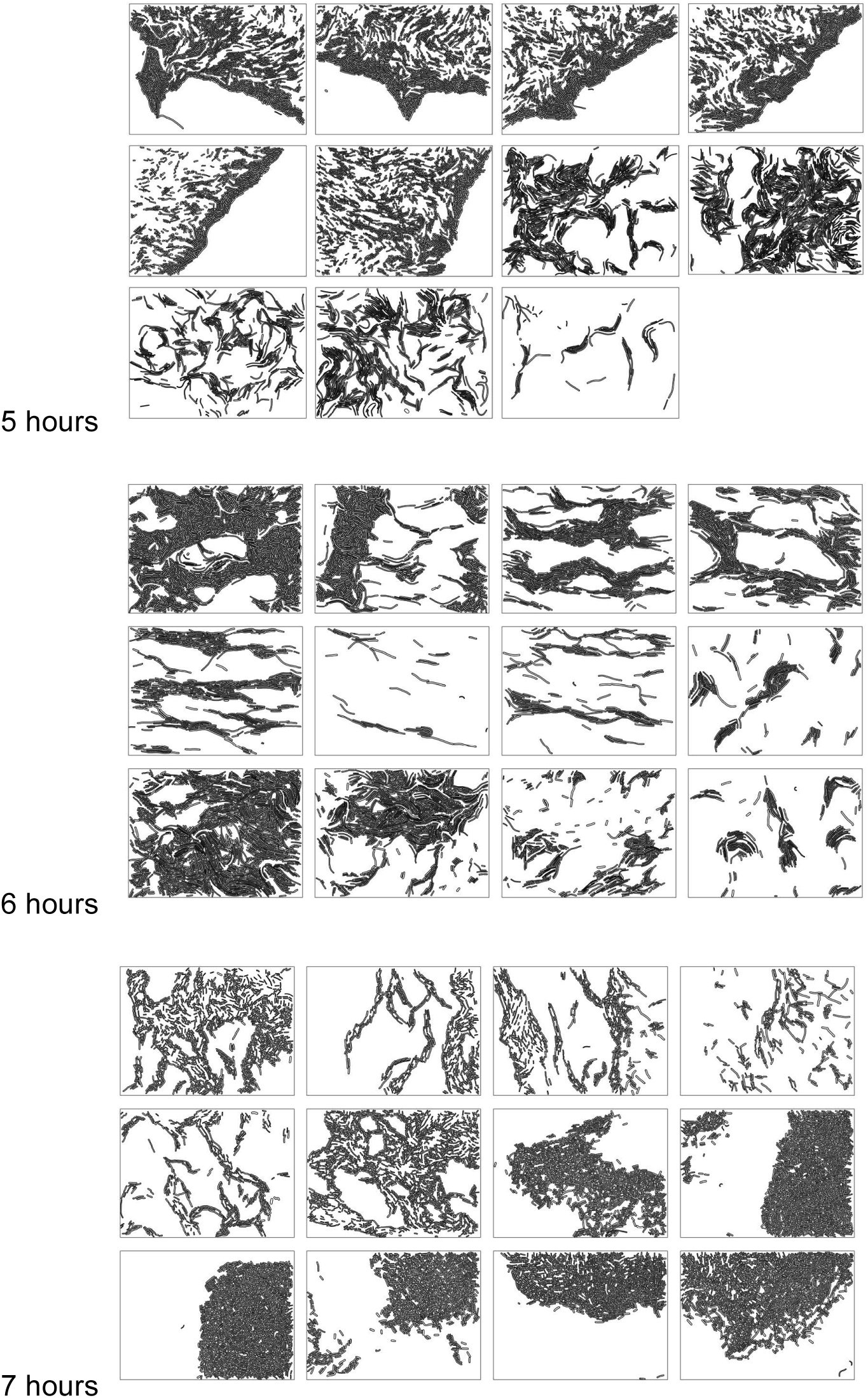
Scatterplots of wild-type *P. mirabilis* strain BB2000 used for analysis. Each individual image is ∼ 126.4 µm wide by ∼ 94.4 µm tall. Images correspond to the phase contrast micrograph in the previous figure, assembled in the same layout.

**Supplemental Figure 4.**
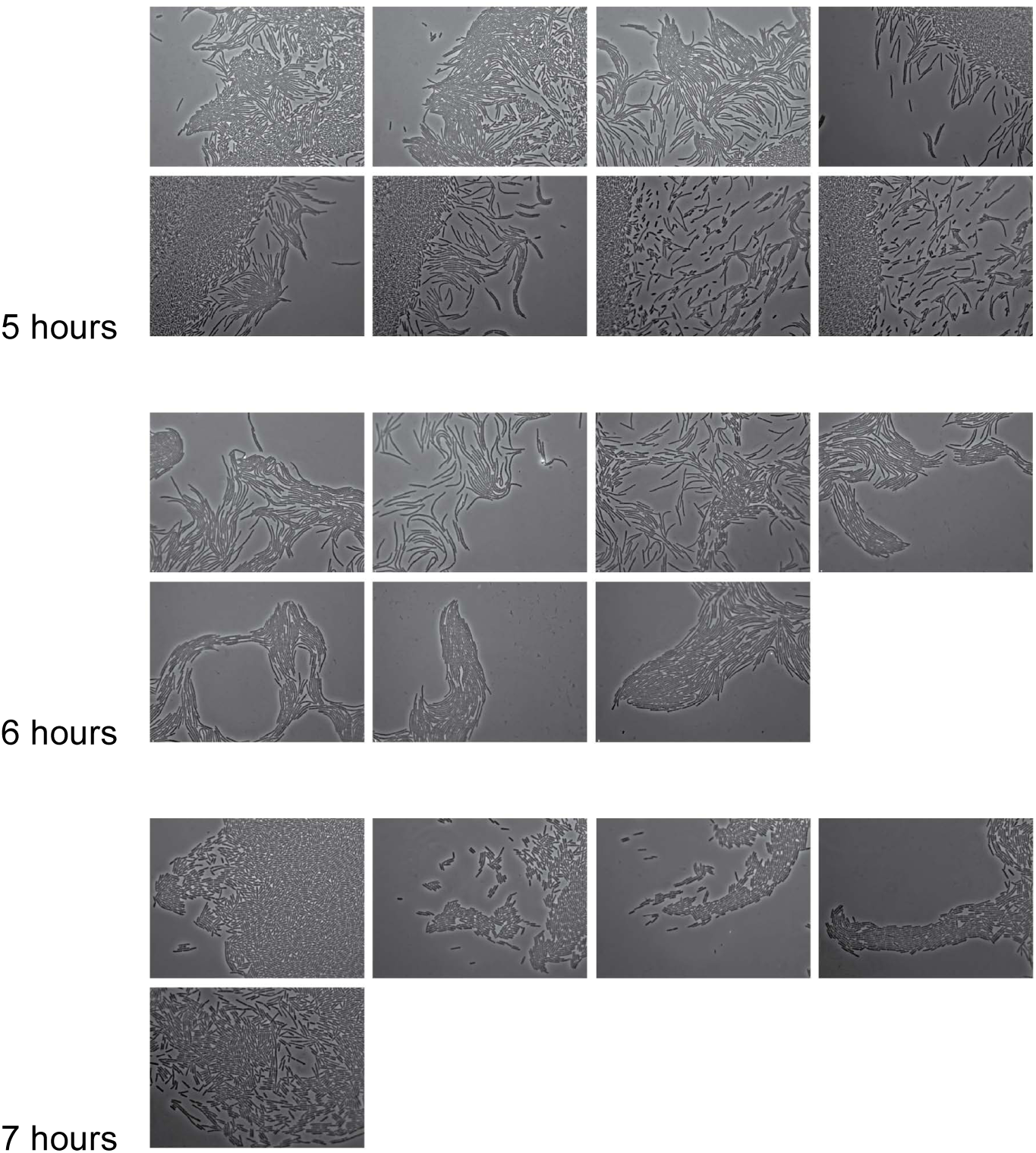
Phase contrast images of the *Δids* strain used for analysis. Each individual image is ∼ 126.4 µm wide by ∼ 94.4 µm tall.

**Supplemental Figure 5.**
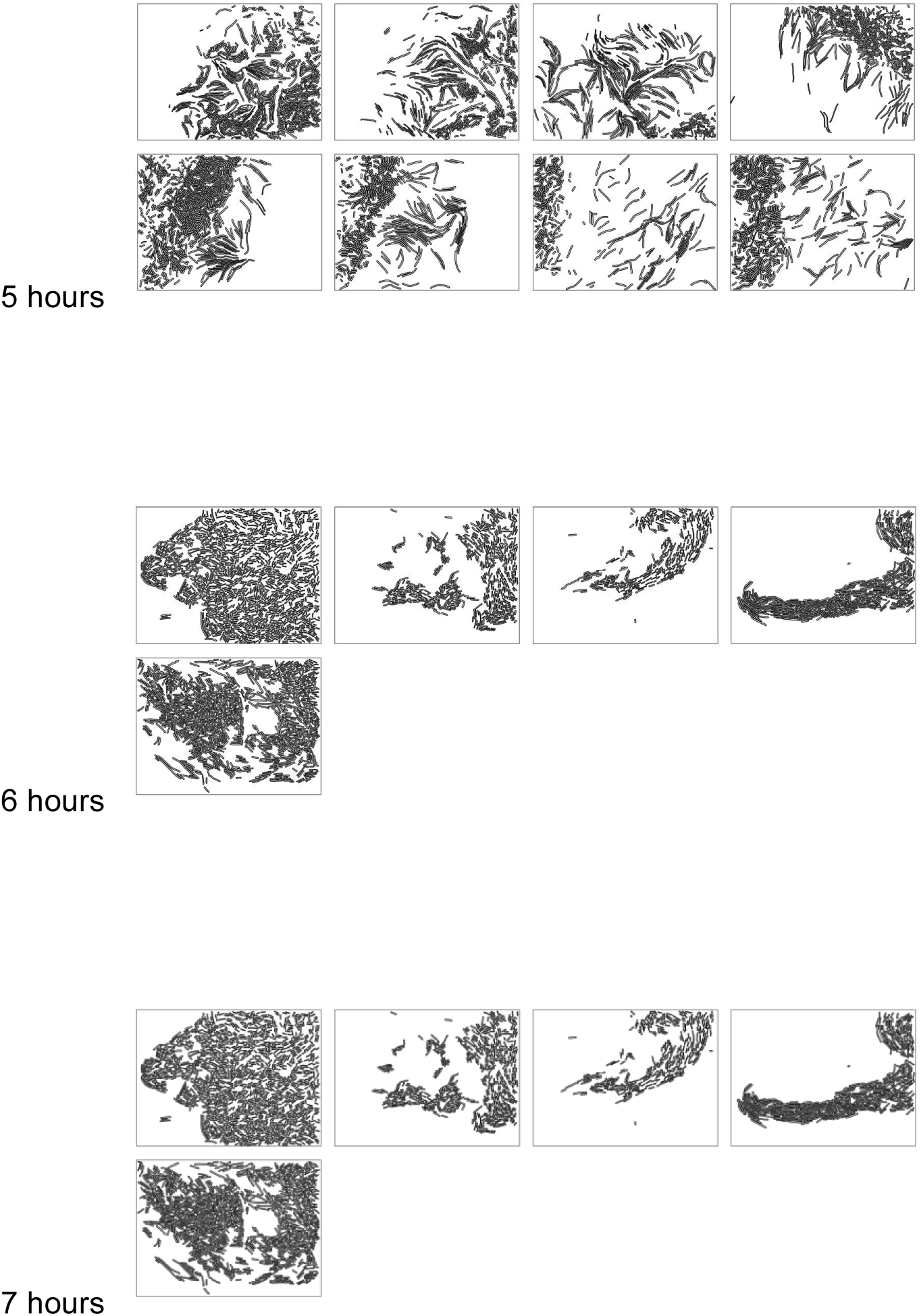
Scatterplots of the *Δids* strain used for analysis. Each individual image is ∼ 126.4 µm wide by ∼ 94.4 µm tall. Images correspond to the phase contrast micrograph in the previous figure, assembled in the same layout.

**Supplemental Figure 6.**
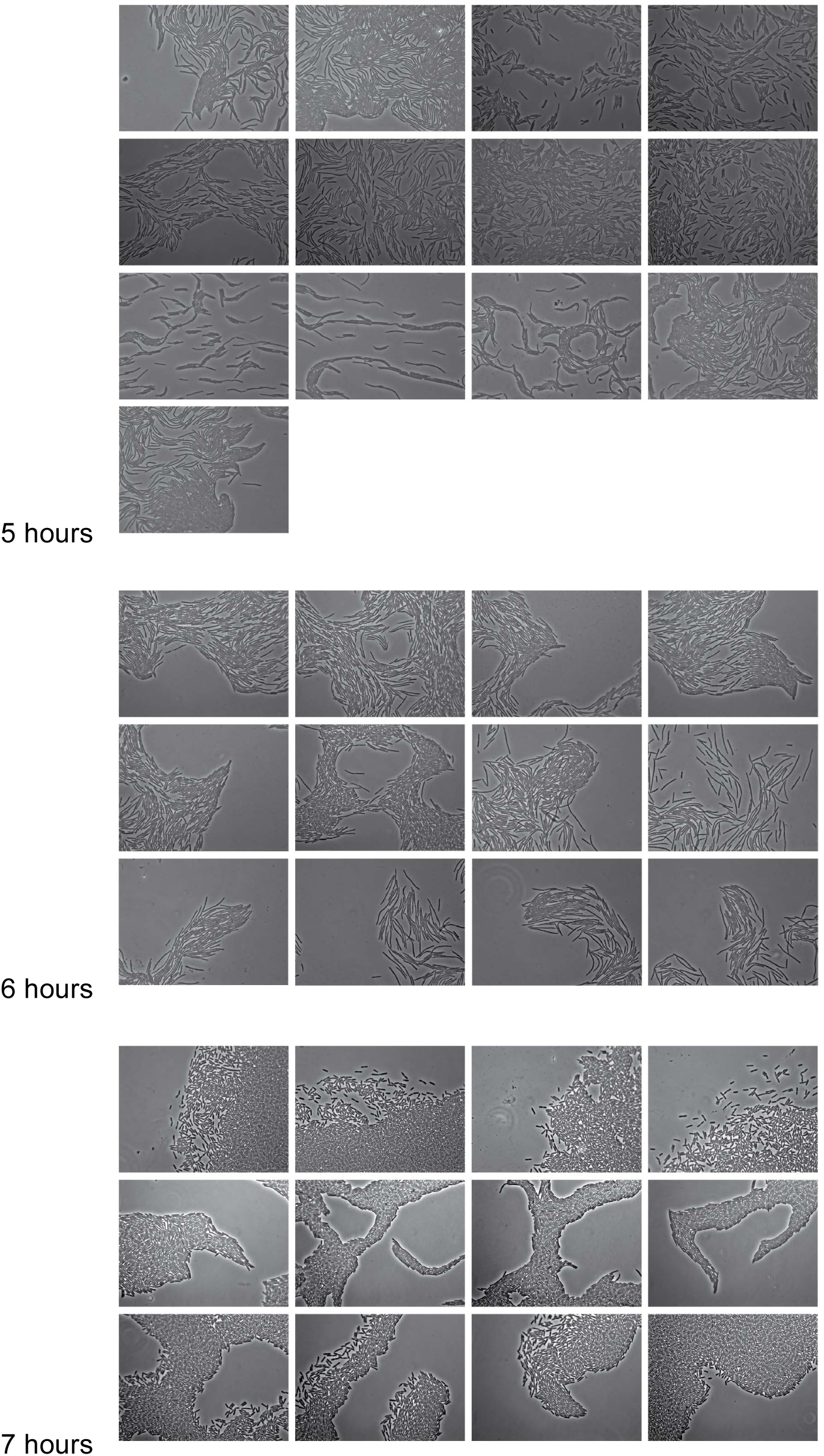
Phase contrast images of the T6S- strain used for analysis. Each individual image is ∼ 126.4 µm wide by ∼ 94.4 µm tall.

**Supplemental Figure 7.**
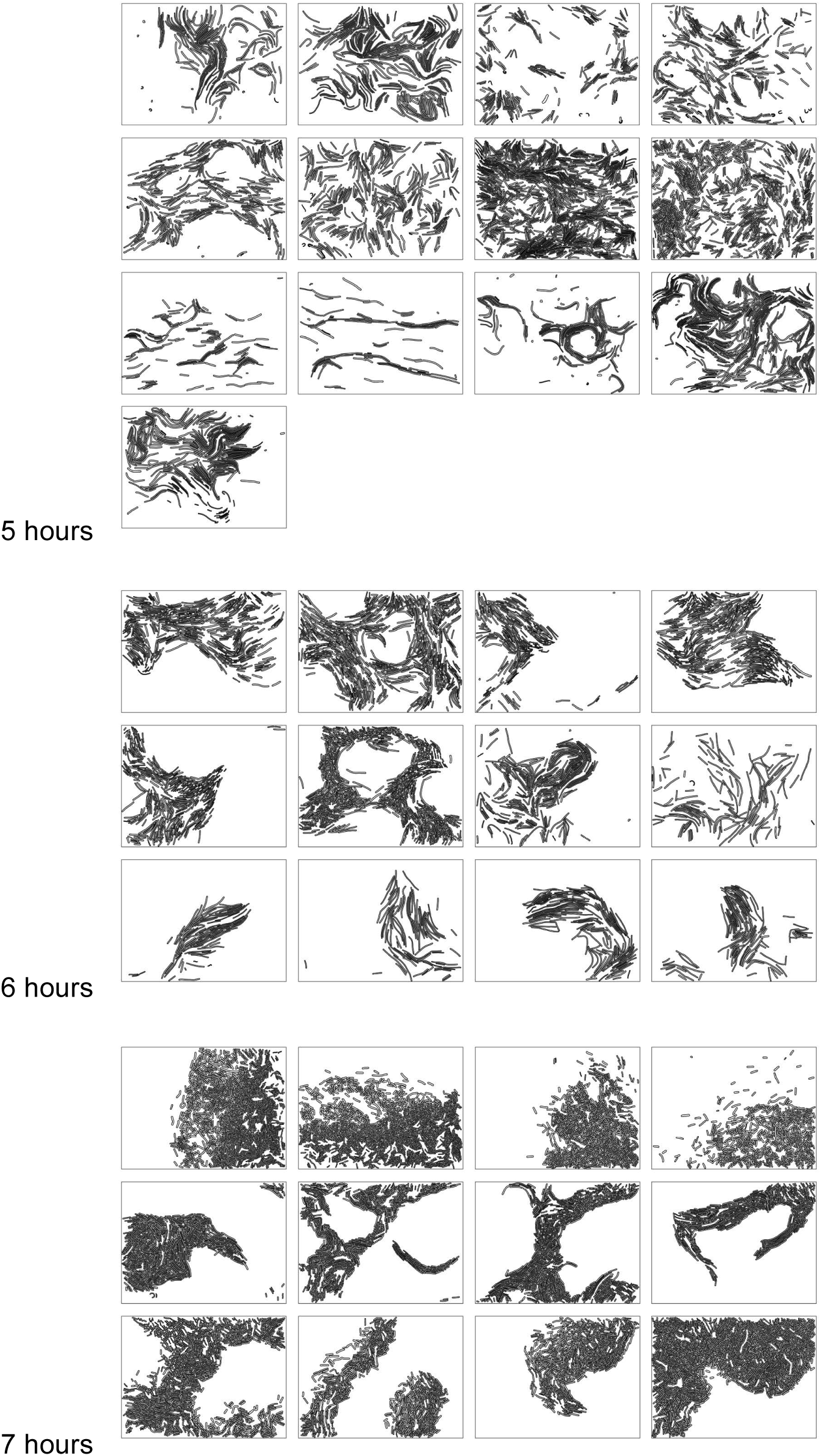
Scatterplots of the T6S- strain used for analysis. Each individual image is ∼ 126.4 µm wide by ∼ 94.4 µm tall. Images correspond to the phase contrast micrograph in the previous figure, assembled in the same layout.

**Supplemental Table 1.** Descriptive statistics for cell lengths for all strains at the measured timepoints.

| Strain | Time | Count | Mean length (µm) | Standard deviation (µm) | Median length (µm) | Interquartile range (µm) | Min (µm) | Max (µm) |
| --- | --- | --- | --- | --- | --- | --- | --- | --- |
| BB2000 | 5 | 8,258 | 4.9 | 3.3 | 3.7 | 3.0 | 2.0 | 36.5 |
| BB2000 | 6 | 7,320 | 5.7 | 3.9 | 4.4 | 3.1 | 2.0 | 38.7 |
| BB2000 | 7 | 10,275 | 3.6 | 1.5 | 3.3 | 1.5 | 2.0 | 36.5 |
| *Δids* | 5 | 6,082 | 5.6 | 4.2 | 3.7 | 4.3 | 2.0 | 35.7 |
| *Δids* | 6 | 1,299 | 12.5 | 6.3 | 11.8 | 8.3 | 2.1 | 45.9 |
| *Δids* | 7 | 4,553 | 4.4 | 2.4 | 3.8 | 2.0 | 2.0 | 39.9 |
| T6S- | 5 | 4,108 | 9.7 | 4.9 | 8.5 | 5.4 | 2.1 | 41.5 |
| T6S- | 6 | 3,949 | 8.1 | 5.4 | 6.4 | 4.9 | 2.0 | 51.6 |
| T6S- | 7 | 9,860 | 4.0 | 1.8 | 3.6 | 1.7 | 2.0 | 30.0 |

**Supplemental Table 2.** The relative distribution of the total populations by hour for cells with lengths over 4 µm or 4 µm and shorter.

| Strain | Time | Percent of cells 4 $\mu\text{m}$ and shorter | Percent of cells longer than 4 $\mu\text{m}$ |
| --- | --- | --- | --- |
| BB2000 | 5 | 55.4 | 44.6 |
| BB2000 | 6 | 42.4 | 57.6 |
| BB2000 | 7 | 72.5 | 27.5 |
| $\Delta ids$ | 5 | 54.6 | 45.4 |
| $\Delta ids$ | 6 | 7.3 | 92.7 |
| $\Delta ids$ | 7 | 56.2 | 43.8 |
| T6S- | 5 | 6.1 | 93.9 |
| T6S- | 6 | 17.0 | 83.0 |
| T6S- | 7 | 61.5 | 38.5 |

